# Targeting CD25-positive lymphoma cells with the antibody-drug conjugate camidanlumab tesirine as single agent or in combination with targeted agents

**DOI:** 10.1101/2023.07.02.547392

**Authors:** Filippo Spriano, Chiara Tarantelli, Luciano Cascione, Eugenio Gaudio, Gaetanina Golino, Lorenzo Scalise, Emanuele Zucca, Anastasios Stathis, Patrick H. Van Berkel, Francesca Zammarchi, Francesco Bertoni

## Abstract

**Introduction:** Camidanlumab tesirine (ADCT-301) is a CD25 specific antibody-drug conjugate (ADC) employing SG3199, a highly cytotoxic DNA minor groove cross-linking pyrrolobenzodiazepine dimer. Camidanlumab tesirine has shown early clinical anti-tumor activity in various cancer types, including B- and T-cell lymphomas. Here, we assessed its preclinical activity as single agent in 57 lymphoma cell lines and in combination with selected drugs in T cell lymphomas-derived cell lines.

**Methods:** Cell lines were exposed to increasing concentrations of camidanlumab tesirine or to SG3199 for 96h followed by MTT proliferation assay. CD25 expression was measured both at cell surface level via fluorescence quantitation and at RNA level, using various technologies. Combination studies were performed exposing cells to increasing doses of camidanlumab tesirine and of additional drugs.

**Results:** Camidanlumab tesirine presented much stronger single agent *in vitro* cytotoxic activity in T than B cell lymphomas. *In vitro* activity was highly correlated with CD25 expression both at cell surface level and RNA level. Based on the higher activity in T cell lymphomas, camidanlumab tesirine-containing combinations were evaluated in cell lines derived from peripheral T cell lymphoma, ALK-pos or ALK-neg anaplastic large cell lymphoma. The most active combination partners were everolimus, copanlisib, venetoclax, vorinostat and pralatrexate, followed by bortezomib, romidepsin, bendamustine and 5-azacytidine.

**Conclusion:** The strong camidanlumab tesirine single agent anti-lymphoma activity and the observed *in vitro* synergisms with targeted agents support further clinical development of camidanlumab tesirine and identify potential combination partners for future clinical studies.

## INTRODUCTION

Antibody-drug conjugates (ADCs) allow the delivery of potent cytotoxic agents to tumor cells and the recent regulatory approval of various ADCs for patients with hematological cancers or solid tumors demonstrates their clinical relevance (1-3). A major factor that contributes to their therapeutic window is the choice of the target (2,3).

The IL-2 receptor (IL2R) exists in two functional forms (4). While the low-affinity form is a dimer, made of a β-(CD122) and a γ-chain (CD132), the high-affinity receptor is a trimeric complex, also including an alpha chain (CD25) (4). High levels of the high-affinity IL2R are transiently expressed by CD4+ and CD8+ T cells following T cell receptor (TCR) activation (5). Unlike other T cell subsets, most regulatory T cells (Tregs) express high levels of CD25 and lower levels of CD122 and CD132. CD25 is also expressed by different types of hematological cancers. All human adult T cell leukemia/lymphoma and hairy cell leukemia constitutively express CD25 (6,7). In addition, CD25 expression has been observed in chronic lymphocytic leukemia (CLL), acute lymphoblastic leukemia, acute myeloid leukemia (AML), diffuse large B cell lymphoma (DLBCL), follicular lymphoma, Hodgkin lymphoma (HL) and peripheral T cell lymphoma (PTCL) (8-10). Due to its pattern of expression, CD25 represents a therapeutic target for antibody and cellular therapy-based approaches in oncology and in autoimmune disorders (11-13).

Camidanlumab tesirine, previously known as ADCT-301, is an ADC composed of the human IgG1 HuMax-TAC directed against CD25, stochastically conjugated through a protease cleavable dipeptide linker to a pyrrolobenzodiazepine (PBD) dimer warhead (SG3199) (14). Upon binding to CD25, camidanlumab tesirine internalizes and traffics to the lysosomes where the PBD dimers are released which form highly cytotoxic DNA interstrand cross-links causing cell death (14).

Phase 1 and phase 2 studies have been conducted exploring camidanlumab tesirine in advanced solid tumors, AML and in relapsed refractory Hodgkin and non-Hodgkin lymphoma (15-17). In the phase 1 study, conducted in relapsed or refractory (R/R) classical Hodgkin lymphoma (cHL) and non-Hodgkin lymphoma, 133 patients were enrolled (77 classical HL and 56 non-Hodgkin lymphoma). Dose-limiting toxicities were reported in five (6%) of 86 patients who were evaluable and grade 3 or worse treatment-emergent adverse events were reported in around 10% of 133 patients. Notably, anti-tumoral activity was seen in classical HL and non-Hodgkin lymphomas (16). In a phase 2 trial in patients with R/R cHL, that had received ≥3 prior systemic therapies including brentuximab vedotin and anti-PD-1, camidanlumab tesirine showed an overall response rate of 70%, while 33% of patients had complete response (18).

The aim of this study was to assess the preclinical activity of camidanlumab tesirine as single agent in a large collection of lymphoma cell lines and its potential role as combination partner.

## METHODS

### Cell lines

Lymphoma cell lines were cultured according to the recommended conditions, as previously described (19). All media were supplemented with fetal bovine serum (10% or 20%) and penicillin-streptomycin-neomycin (≈5,000 units penicillin, 5 mg streptomycin, and 10 mg neomycin/mL; Sigma). Human cell line identities were confirmed by short tandem repeat DNA fingerprinting using the Promega GenePrint 10 System kit (B9510). Cells were periodically tested for mycoplasma negativity using the MycoAlert Mycoplasma Detection Kit (Lonza).

### Cytotoxic activity in single and combination

Anti-proliferative activity after treatment was assessed as previously described (20). Briefly, cells were exposed to ADCT-301, B12-SG3249 or to SG3199 for 96h and assayed by MTT [3-(4,5-dimethylthiazolyl-2)-2, 5-diphenyltetrazoliumbromide]. Synergism assessment was done exposing cells (96h) to increasing doses of camidanlumab tesirine and of other agents alone or in combination, followed by MTT assay, and determination of the Chou-Talalay combination index (CI) concentrations, as previously described (20,21). Concentrations of compounds that after 96h of treatment left 10% or less of proliferating cells already with the single agent were discarded from further analyses. Combinations were defined as synergistic (median CI < 0.9), additive (median CI, 0-9-1.1) or of no benefit/antagonist (median CI > 1.1).

### Compounds

Camidanlumab tesirine, SG3199 (warhead) and B12-SG3249 (isotype control ADC) were provided by ADC Therapeutics. Everolimus, pralatrexate, vorinostat, bortezomib, venetoclax, copanlisib, romidepsin, bendamustine, and 5-azacytidine were purchased from Selleckchem (Houston, TX, USA).

### CD25 expression

The Quantum Simply Cellular (QSC) anti-Human IgG beads from Bangs Laboratories were utilized to establish a calibration curve for determining the absolute CD25 surface expression. Subsequently, the Antibody Binding Capacity (ABC) values were normalized to the control isotype antibody B12.

CD25 RNA expression values were extracted from previously reported datasets obtained using Illumina HT-12 arrays (GSE94669 (21)), HTG EdgeSeq Oncology Biomarker Panel (GSE103934 (21)) and total-RNA-Seq (GSE221770 (19)).

### Data Analysis

Pearson correlation (r) was calculated for IC50 values versus cell surface CD25 expression levels and vs RNA levels and for all the other correlations. The presence of *BCL2* and/or *MYC* translocations and *TP53* inactivation were retrieved from our previous publication(21). Differences in IC50 values among lymphoma subtypes were calculated using the non-parametric Mann-Whitney t-test. Statistical significance was defined by P values of 0.05 or less. Statistical analyses, correlations and boxplots were performed using GraphPad Prism.

## RESULTS

### The *in vitro* anti-tumor activity of camidanlumab tesirine is dependent on CD25 expression

A panel of lymphoma cell lines were exposed to increasing concentrations of the CD25 targeting ADC camidanlumab tesirine for 96 hours. The human lymphoma cell lines were derived from activated B cell like (ABC; n.=7) and germinal center B cell like (GCB) DLBCL (n.=19), mantle cell lymphoma (MCL; n.=10), marginal zone lymphoma (MZL; n.=6), ALK+ anaplastic large cell lymphoma (ALCL; n.=4), cutaneous T cell lymphoma (CTCL; n.=4), CLL (n.=2), HL (n.=3), primary mediastinal B cell lymphoma and PTCL (n.=1).

The median IC50 of camidanlumab tesirine was 650 pM across all cell lines (Table 1, Supplementary Table 1). The isotype control ADC B12-SG3249, on the contrary, did not show activity in cell lines sensitive to camidanlumab tesirine (Supplementary Table 1). The cytotoxic activity of camidanlumab tesirine was highly dependent on CD25 expression as shown by a negative correlation between IC50 values and both CD25 protein levels on cell surface (n=57, Pearson r = -0.369, P = 0.0047) as well as CD25 RNA levels (n=50, Pearson r = -0.56, P<0.0001 (Illumina arrays); n=36, Pearson r = -0.59, P<0.0001 (HTG); n=53, Pearson r = - 0.65, P<0.0001 (total RNA-seq) (Figure 1). To further underline this association, cell lines could be divided in two groups based on their CD25 expression and sensitivity to camidanlumab tesirine. Cells having IC50 below 5 pM had a significantly higher CD25 expression compared to cells with IC50 higher that 5 pM (P<0.0001) (Figure 2).

**Table 1.** Camidanlumab tesirine has *in vitro* anti-proliferative activity in lymphoma cell lines. IC50 value distributions in 60 lymphoma cell lines. MTT proliferation assay and IC50 calculation on cell lines exposed (96h) to increasing camidanlumab tesirine concentrations. DLBCL, diffuse large B cell lymphoma; ABC, activated B cell; GCB, germinal center B cell; MCL, mantle cell lymphoma; MZL, marginal zone lymphoma; CLL, chronic lymphocytic leukemia; HL. Hodgkin’s lymphoma; PMBCL, primary mediastinal large B cell lymphoma; ALCL, anaplastic large cell lymphoma; CTCL, cutaneous T cell lymphoma; PTCL, peripheral T cell lymphoma-not otherwise specified. n/a, not determined.

| Histotype | Number of cell lines | Median IC50 (pM) | 95% CI of median (pM) |
| --- | --- | --- | --- |
| ABC DLBCL | 7 | 750 | 300-6,500 |
| GCB DLBCL | 19 | 1250 | 6.3-4,000 |
| MCL | 10 | 825 | 400-2,750 |
| MZL | 6 | 75.63 | 0.225- 800 |
| PTCL | 1 | 3 | n/a |
| PMBCL | 1 | 2 | n/a. |
| HL | 3 | 3,500 | 350-20,000 |
| ALK+ ALCL | 4 | 3.5 | 1.45-8.5 |
| CLL | 2 | 2,629 | 7.25-5,250 |
| CTCL | 4 | 845 | 1.5-40,000 |
| Murine B cell lymphoma | 2 | 1,475 | 500-2,450 |
| Canine B cell lymphoma | 1 | 225 | n/a |

**Figure 1.**
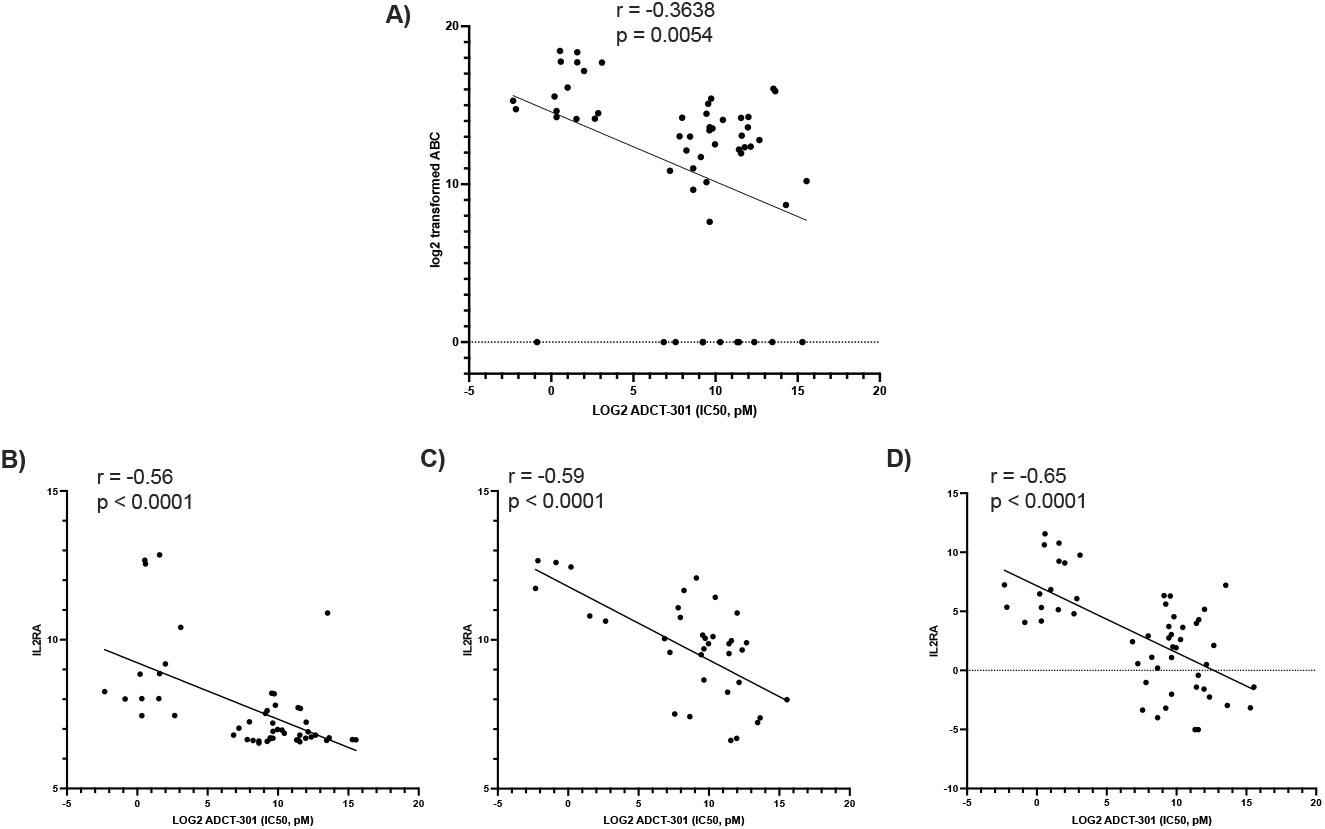
Correlation between camidanlumab tesirine (ADCT-301) IC50 and CD25 protein surface or RNA expression. A) Correlation between camidanlumab tesirine (ADCT-301) IC50 and CD25 surface expression. B), C), D) Correlation between camidanlumab tesirine (ADCT-301) IC50 and CD25 (*IL2RA*) RNA expression analyzed by Illumina arrays, HTG or Total RNA-seq respectively. Pearson correlation performed. ABC = Antibody binding capacity.

**Figure 2.**
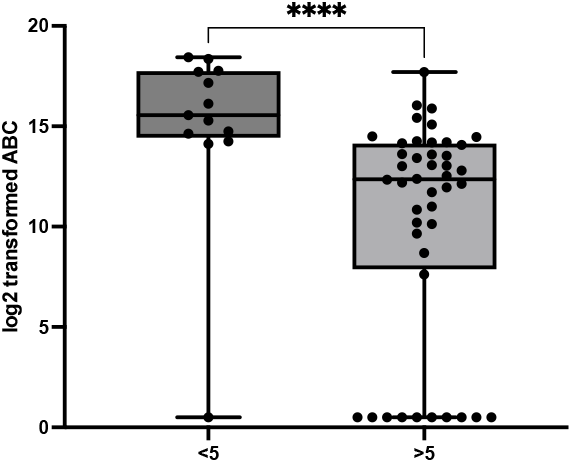
CD25 surface expression in cell lines where camidanlumab tesirine has IC50 higher or lower than 5 pM. ABC = Antibody binding capacity. Mann-Whiney test performed; **** = p<0.0001

Among the individual lymphoma histotypes, the most sensitive cell lines were those derived from ALCL and PTCL (Table 1), which were also the two subtypes with the highest CD25 surface expression (Figure 2). Indeed, CD25 surface expression was much higher in T (n=9, median antibody binding capacity 21,2921; 95% C.I., 10,915-334,885) than in B cell lymphomas (n=48, median antibody binding capacity 6,511 95% C.I., 2,055-17,910) (P = 0.0024) (Figure 3). This difference was matched with the stronger cytotoxic activity of camidanlumab tesirine in cell lines derived from T cell (n=6, median IC50=3 pM; 95% C.I., 1.5 pM-0.9 nM) than from B cell lymphomas (n=48, 725 pM; 95% C.I., 350 pM-2.5 nM) (P = 0.0024) (Figure 3).

**Figure 3.**
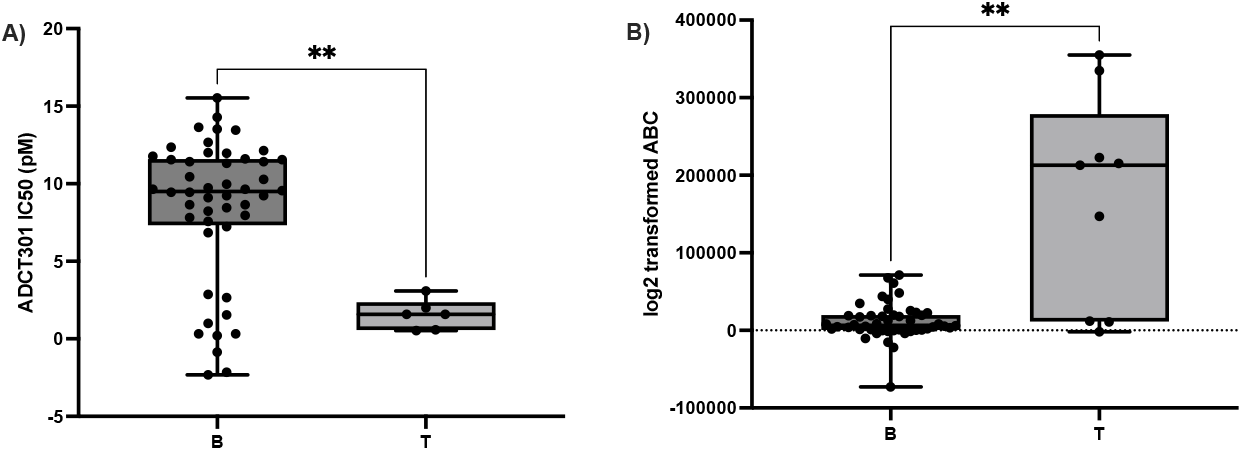
Sensitivity to camidanlumab tesirine (ADCT-301) and CD25 expression in B vs T cell lymphoma. A) Camidanlumab tesirine (ADCT-301) IC50 (pM) in B cell lymphoma compared to T cell lymphoma cell lines. B) CD25 surface expression in B cell lymphoma compared to T cell lymphoma cell lines. ABC = Antibody binding capacity. Mann-Whiney test performed; ns = non-significant; ** = p<0.01

Camidanlumab tesirine was also tested in three non-human lymphoma cell lines. IC50 values were 2.5 nM and 500 pM in two mouse cell lines, and 225 pM in a canine DLBCL cell line (Supplementary Table 1), indicating that camidanlumab tesirine is not cross-reactive with mouse and dog CD25. Indeed a mouse CD25 cross-reactive surrogate ADC for camidanlumab tesirine has been develop to perform studies with the ADC in syngeneic mouse models (22).

Within the group of DLBCL cell lines, there was no association between sensitivity to camidanlumab tesirine and the presence of *BCL2* and *MYC* translocations or *TP53* inactivation (Figure S1), similarly reported for other ADCs containing new generations of DNA targeting payloads (23,24).

### The *in vitro* cytotoxic activity of camidanlumab tesirine is correlated with its warhead

In parallel to the ADC, all the cell lines were also exposed to SG3199, camidanlumab tesirine’s warhead. (Table S1) When we correlated the IC50 obtained with camidanlumab tesirine against the anti-tumor activity of SG3199, cell lines appeared divided in two clusters, driven by the sensitivity to camidanlumab tesirine in line with what observed regarding CD25 expression (Figure 4.). For both the camidanlumab tesirine sensitive and the camidanlumab tesirine resistant cell lines, the pattern of activity of the ADC was correlated to its warhead. The correlations were weaker among the sensitive (SG3199; r = 0.62, P = 0.01), while they appeared stronger in the resistant group (SG3199, r = 0.8, P < 0.0001) (Figure 4). Differently from camidanlumab tesirine, SG3199 was active also in the murine and canine cell lines (Table S1).

**Figure 4.**
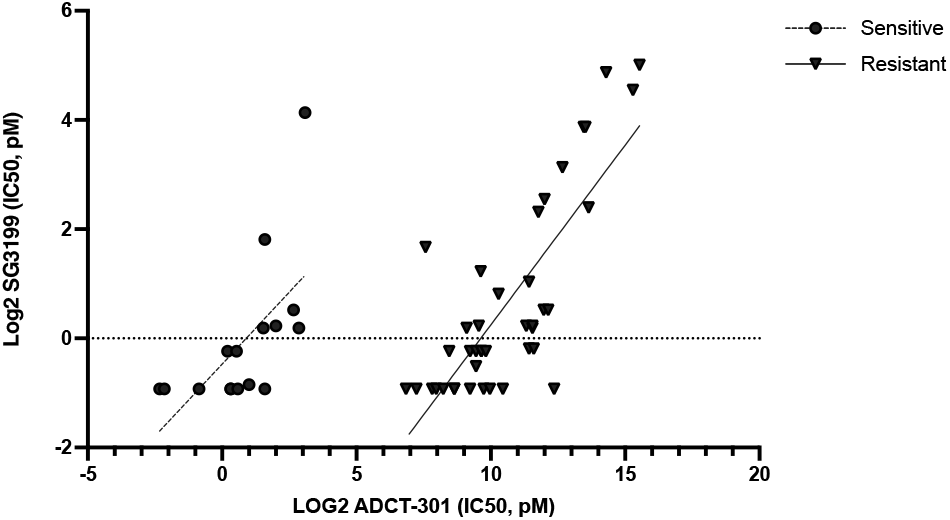
Correlation between IC50 values of camidanlumab tesirine (ADCT-301) and its warhead SG3199.

### Camidanlumab tesirine-based combinations are active in T cell lymphoma models

We next explored camidanlumab tesirine-based combinations in T cell lymphoma models. Four cell lines derived from ALK+ ALCL (Karpas-299, KI-JK), CTCL (MAC-1) and PTCL-NOS (FE-PD) were exposed to camidanlumab tesirine in combination with drugs known to have clinical activity against T cell lymphomas (25) (Table 2). Camidanlumab tesirine plus the mTOR inhibitor everolimus showed synergism in all four cell lines tested. The combination with the folate antagonist pralatrexate was synergistic in the two ALK+ ALCL cell lines tested. Combination with the class I and II HDAC inhibitor vorinostat was synergistic in three cell lines (CTCL, ALK-, ALK+ ALCL) and additive in the remaining ALK+ ALCL (cell line Karpas-299). The combination with bortezomib led to synergism in the CTCL and ALK-ALCL cells and additivity in the two ALK+ ALCL cells. Combinations with the BCL2 inhibitor venetoclax was synergistic in both ALK+ALCL and in the CTCL but was not beneficial in the PTCL-NOS. The PI3K inhibitor copanlisib synergized with camidanlumab tesirine in the CTCL, the ALK-ALCL and in only one of the two ALK+ALCL (Karpas-299). cell lines. The latter two also achieved synergy with the addition of the class I and II HDAC inhibitor romidepsin, while no benefit was seen in the other cell lines. The combination with the chemotherapy agent bendamustine showed synergy in one ALK+ ALCL (Karpas-299) and in the ALK-ALCL, but no benefit in the two other models. Finally, the demethylating agent 5-azacytidine was synergistic only in one ALK+ ALCL (Karpas-299) and it was of no benefit in the remaining three cell lines (Table 2).

**Table 2.** Camidanlumab tesirine containing combinations in T cell lymphoma cell lines. 95% C.I.: 95% confidence interval; PTCL-NOS, peripheral T cell lymphoma-not otherwise specified; ALCL, anaplastic large cell lymphoma; CTCL, cutaneous T cell lymphoma. Synergism, additive and no benefit effects were defined using the Chou-Talalay Combination Index (CI). Synergism, CI < 0.9; additive effect, 0.9 < CI < 1.1; no benefit, CI >1.1.

| Combination Partner | Histology | Cell line | Median combination index | 95% C.I. |
| --- | --- | --- | --- | --- |
| Bendamustine | PTCL-NOS | FE-PD | 0.91 | 0.45 - 1.75 |
|  | ALK+ALCL | Karpas-299 | 0.57 | 0.29 - 0.84 |
|  | ALK+ALCL | KI-JK | 1.17 | 0.57 - 2.42 |
|  | CTCL | MAC1 | 1.73 | 0.33 - 5.71 |
| Copanlisib | PTCL-NOS | FE-PD | 0.66 | 0.22 - 1.47 |
|  | ALK+ALCL | Karpas-299 | 0.84 | 0.47 - 2.31 |
|  | ALK+ALCL | KI-JK | 1.26 | 0.84 - 1.9 |
|  | CTCL | MAC1 | 0.38 | 0.17 - 0.96 |
| Venetoclax | PTCL-NOS | FE-PD | 1.68 | 0.67 - 2.1 |
|  | ALK+ALCL | Karpas-299 | 0.44 | 0.27 - 0.68 |
|  | ALK+ALCL | KI-JK | 0.55 | 0.38 - 0.67 |
|  | CTCL | MAC1 | 0.42 | 0.32 - 0.57 |
| Everolimus | PTCL-NOS | FE-PD | 0.05 | 0.03 - 0.07 |
|  | ALK+ALCL | Karpas-299 | 0.03 | 0.02 - 0.04 |
|  | ALK+ALCL | KI-JK | 0.03 | 0.02 - 0.05 |
|  | CTCL | MAC1 | 0.74 | 0.27 - 1.18 |
| 5-azacytidine | PTCL-NOS | FE-PD | >3 | >3 |
|  | ALK+ALCL | Karpas-299 | 0.6 | 0.36 - 0.97 |
|  | ALK+ALCL | KI-JK | >3 | 0.38 - >3 |
|  | CTCL | MAC1 | 1.87 | 0.70 - >3 |
| Bortezomib | PTCL-NOS | FE-PD | 0.13 | 0.06 - 0.1985 |
|  | ALK+ALCL | Karpas-299 | 0.93 | 0.4412 - 1.31 |
|  | ALK+ALCL | KI-JK | 1.03 | 0.62 - 1.73 |
|  | CTCL | MAC1 | 0.31 | 0.17 - 0.41 |
| Pralatrexate | ALK+ALCL | Karpas-299 | 0.5 | 0.19 - 1.44 |
|  | ALK+ALCL | KI-JK | 0.5 | 0.18 - 1.28 |
| Romidepsin | PTCL-NOS | FE-PD | 1.16 | 0.81 - 8.3 |
|  | ALK+ALCL | Karpas-299 | 0.85 | 0.64 - 1.1 |
|  | ALK+ALCL | KI-JK | 0.89 | 0.79 - 1.19 |
|  | CTCL | MAC1 | 1.25 | 0.43 - >3 |
| Vorinostat | PTCL-NOS | FE-PD | 0.85 | 0.64 - 2.08 |
|  | ALK+ALCL | Karpas-299 | 0.96 | 0.43 - 1.44 |
|  | ALK+ALCL | KI-JK | 0.55 | 0.47 - 0.59 |
|  | CTCL | MAC1 | 0.4 | 0.11 - 0.72 |

## DISCUSSION

In this manuscript, we show that camidanlumab tesirine has strong *in vitro* cytotoxic activity in models of CD25-positive lymphomas, and its anti-tumor activity increases when combined with a series of additional anti-cancer agents.

Camidanlumab tesirine has been initially studied in four CD25-positive cell lines (two ALCL, two HL) and three CD25-negative cell lines (two Burkitt lymphoma, one CTCL) (14). In this publication, camidanlumab tesirine had cytotoxic activity only in the CD25-positive cell lines, and no correlation between expression and activity among the positive cell lines could be established (14). Here, we extended the analysis to a much wider panel of cell lines derived from both B and T cell mature lymphomas with different CD25 expression levels. Increasing the number of cell lines led to a strong association between CD25 expression and camidanlumab tesirine anti-tumor activity. The anti-tumor activity was highly dependent on CD25 expression, both at protein level on the cell surface, and, even more at RNA level, independently from the technologies employed to measure it. Noteworthy, the latter also included a targeted RNA-Seq approach specifically designed for formalin fixed paraffin embedded materials, thus, easily transferrable to clinical specimens. The higher correlation with CD25 RNA levels might be due to technical issues that make the flow cytometry measurements to assess CD25 protein levels less robust than RNA-based techniques.

In line with the pattern of CD25 expression in clinical specimens (6-10), we observed a higher expression of CD25 in cell lines derived from mature T cell lymphomas. This was reflected by a higher *in vitro* camidanlumab tesirine anti-tumor activity in T than B cell lymphomas. This finding sustains what was observed in the phase 1 study, in which the ORR in patients with T cell lymphoma was twice compared to what observed in the B cell lymphoma population (48% vs 23%) (16). In the phase 1 study, the highest ORR and CR rates were seen in r/r HL (71% and 42%, respectively) (16), which was later confirmed in the phase 2 study (70% and 33%, %, respectively) (18). Interestingly, the three HL-derived cell lines tested in our study, and which differed from the ones previously tested (14), were not sensitive to camidanlumab tesirine. Of interest, in the phase 1 study, a higher CD25 histoscore was associated with higher ORR among HL patients (16). A correlation between the anti-tumor activity of an ADC and its target expression is often, but not always, observed due to complex interaction between characteristics of the ADC and of its target in the tumor cells (24,26-33).

Both among the camidanlumab tesirine sensitive and the camidanlumab tesirine resistant cell lines, the pattern of activity of the ADC was correlated to that of its warhead SG3199. The correlation was weaker among the sensitive models, while it appeared stronger in the resistant group. These data suggest that the strongest factors to drive sensitivity to camidanlumab tesirine are CD25 expression levels and the intrinsic degree of sensitivity of each tumor to the PBD dimer warhead SG3199. The latter point highlights the importance of optimizing the warheads as a fundamental step to fully exploit the potential offered by the targeted delivery that an ADC facilitates.

Finally, focusing on the T cell lymphoma models, we explored the potential benefit of adding camidanlumab tesirine to drugs with antitumor activity reported in T cell lymphomas (34-36). We observed synergism especially when we combined camidanlumab tesirine with everolimus, vorinostat, bortezomib, copanlisib, venetoclax and pralatrexate.

The combination with the mTOR inhibitor everolimus appeared very active, albeit its move to the clinical setting might be difficult based on the toxicity observed in a phase 1 study that explored the combination of another mTOR inhibitor, temsirolimus, with inotuzumab ozogamicin, a CD22 targeting ADC also containing a DNA damaging toxin, i. e. a semi-synthetic derivative of calicheamicin (37).

Conversely, the CD33 targeting gemtuzumab ozogamicin, containing the same payload as inotuzumab ozogamicin, has been safely combined with class I and II HDAC inhibitors in AML patients, indicating that the combination of camidanlumab tesirine with this type of epigenetic agents could be feasible in patients with T cell lymphomas, in which they are approved as single agents.

PI3K inhibitors are potentially interesting agents for T cell lymphoma patients (35,38). Our data shows the potential benefit of combining them with camidanlumab tesirine. Furthermore, recent data indicate that the direct anti-tumor activity of this combination might also be boosted by the expected depletion of the tumor infiltrating Tregs, which express CD25 and depend on PIKδ (19,22,39).

Pralatrexate has been safely combined with standard CHOP (Fol-CHOP) in a phase 1 study for 1st line PTCL patients, making the combination with camidanlumab tesirine to be considered for clinical evaluation. Venetoclax has shown low clinical activity in PTCL patients (40), but it might work more in combination (36), as observed here with camidanlumab tesirine. Hints on the feasibility of this combination will come from the ongoing phase 1 trial exploring the CD19 targeting ADC loncastuximab tesirine (ADCT-402), based on the same payload as camidanlumab tesirine, and venetoclax for R/R lymphoma patients (NCT05053659).

The results obtained for camidanlumab tesirine in combination with bendamustine are perhaps slightly worse than what has been reported for the combination of camidanlumab tesirine with gemcitabine in three lymphoma models (synergism in in two HL and in one ALCL cell lines) (41). However, benefit has been previously reported for the combination of the BCMA targeting, PBD-containing, MEDI2228 with bortezomib in multiple myeloma models (42), supporting our data with camidanlumab tesirine together with the same proteasome inhibitor. So far, clinical safety data are only available for the combination of bortezomib with the BCMA targeting belantamab mafodotin, which has a tubulin-damaging payload (43).

In conclusion, our results show strong single agent *in vitro* anti-tumor activity for camidanlumab tesirine in CD25 expressing lymphoma models, especially in T cell lymphomas. Our data also identified possible combination partners that could be clinically explored, including HDAC inhibitors, PI3K and BCL2 inhibitors and folate antagonists.

## Acknowledgments

This project was partially supported by research funds from ADC Therapeutics.

## Authors contributions

FS: performed experiments, performed data mining, interpreted data, co-wrote the manuscript.

CT, EG, LC: performed experiments, interpreted data.

LC: performed data mining.

GG, LS: performed experiments.

EZ, AS: provided advice.

PHVB, FZ: co-designed the study, provided reagents, supervised the study.

FB: co-designed the study, performed data mining, interpreted data, supervised the study and co-wrote the manuscript.

All authors reviewed and accepted final version of the manuscript.

