## supplementary table and figure for "Targeting CD25-positive lymphoma cells with the antibody-drug conjugate camidanlumab tesirine as single agent or in combination with targeted agents"

### SUPPLEMENTARY MATERIAL

**Table S1. List of individual camidanlumab tesirine, B12-SG3249 and SG3199 IC50 values, assessed at 96 hours by MTT, and CD25 protein surface expression values, measured by FACS.**

| Histology | Cell line | ADCT-301 (IC50, pM) | CD25 Log2 normalized antibody binding capacity | SG3199 (IC50, pM) | B12-SG3249 (IC50, pM) |
|---|---|---|---|---|---|
| ABC DLBCL | HBL-1 | 2562.5 | 0.0 | 1.2 | 2500.0 |
| ABC DLBCL | OCI-Ly-3 | 600 | 0.0 | 0.8 | 1200.0 |
| ABC DLBCL | RCK8 | 750 | 15.1 | 1.2 | 2500.0 |
| ABC DLBCL | RI-1 | 4100 | 14.3 | 5.8 | 3162.5 |
| ABC DLBCL | TMD8 | 300 | 12.1 | 0.5 | 550.0 |
| ABC DLBCL | U2932 | 6500 | 12.8 | 8.8 | 5562.5 |
| ABC DLBCL | SU-DHL-2 | 550 | 11.7 | 1.1 | 1300.0 |
| GCB DLBCL | DB | 190 | 0.0 | 3.2 | 300.0 |
| GCB DLBCL | DOHH2 | 115 | 0.0 | 0.5 | 125.0 |
| GCB DLBCL | FARAGE | 3100 | 13.1 | 0.9 | 1650.0 |
| GCB DLBCL | Karpas-422 | 12750 | 15.9 | 5.3 | 8000.0 |
| GCB DLBCL | OCI-Ly-1 | 6.3 | 14.2 | 1.4 | 275.0 |
| GCB DLBCL | OCI-Ly-18 | 0.6 | 0.0 | 0.5 | 150.0 |
| GCB DLBCL | OCI-Ly-19 | 225 | 13.0 | 0.5 | 175.0 |
| GCB DLBCL | OCI-Ly-7 | 2.9 | 14.1 | 1.1 | 2250.0 |
| GCB DLBCL | OCI-Ly-8 | 0.2 | 15.3 | 0.5 | 90.0 |
| GCB DLBCL | PFEIFFER | 11750 | 16.0 | 14.6 | 25000.0 |
| GCB DLBCL | SU-DHL-10 | 2750 | 0.0 | 0.9 | 750.0 |
| GCB DLBCL | SU-DHL-16 | 3000 | 12.0 | 1.1 | 2500.0 |
| GCB DLBCL | SU-DHL-4 | 4000 | 13.6 | 1.4 | 2250.0 |
| GCB DLBCL | SU-DHL-5 | 700 | 10.1 | 0.8 | 650.0 |
| GCB DLBCL | SU-DHL-6 | 11250 | 0.0 | 14.6 | 9500.0 |
| GCB DLBCL | SU-DHL-8 | 3000 | 14.2 | 1.2 | 2500.0 |
| GCB DLBCL | TOLEDO | 4500 | 12.4 | 1.4 | 2812.5 |
| GCB DLBCL | VAL | 1.3 | 14.6 | 0.5 | 340.0 |
| GCB DLBCL | WSU-DLCL2 | 1250 | 0.0 | 1.8 | 1500.0 |
| MCL | GRANTA519 | 400 | 11.0 | 0.5 | 400.0 |
| MCL | JEKO1 | 1400 | 14.1 | 0.5 | 2250.0 |
| MCL | JVM2 | 2750 | 12.2 | 2.0 | 3000.0 |
| MCL | MAVER1 | 700 | 14.5 | 0.7 | 900.0 |
| MCL | MINO | 250 | 14.2 | 0.5 | 450.0 |
| MCL | REC1 | 47500 | 10.2 | 32.2 | 32500.0 |
| MCL | SP49 | 850 | 15.4 | 0.5 | 700.0 |
| MCL | SP53 | 1000 | 12.5 | 0.5 | 850.0 |
| MCL | UPN1 | 800 | 7.6 | 0.8 | 600.0 |

| | | | | | |
|---|---|---|---|---|---|
| MCL | Z138 | 400 | 9.6 | 0.5 | 300.0 |
| MZL | ESKOL | 1.3 | 14.3 | 0.5 | 650.0 |
| MZL | HAIRM | 800 | 13.6 | 0.8 | 900.0 |
| MZL | HC1 | 150 | 10.8 | 0.5 | 200.0 |
| MZL | Karpas-1718 | 0.2 | 14.7 | 0.5 | 400.0 |
| MZL | SSK41 | 1.2 | 15.6 | 0.8 | 650.0 |
| MZL | VL51 | 600 | 0.0 | 0.5 | 450.0 |
| HL | AMHLH | 350 | 13.0 | 0.8 | 300.0 |
| HL | KMH2 | 3500 | 12.3 | 5.0 | 5250.0 |
| HL | L428 | 20000 | 8.7 | 29.2 | 13750.0 |
| PMBCL | Karpas-1106P | 2 | 16.1 | 0.6 | 690.0 |
| ALCL | Karpas-299 | 8.5 | 17.7 | 17.5 | 12500.0 |
| ALCL | KIJK | 3 | 17.7 | 3.5 | 2962.5 |
| ALCL | L82 | 4 | 17.2 | 1.2 | 5500.0 |
| ALCL | SU-DHL-1 | 1.5 | 18.4 | 0.8 | 600.0 |
| CLL | MEC1 | 5250 | 0.0 | 0.5 | 1800.0 |
| CLL | PCL-12 | 7.3 | 14.5 | 1.1 | 2562.5 |
| CTCL | MAC1 | 1.5 | 17.8 | 0.5 | 1500.0 |
| CTCL | H9 | 790 | 13.4 | 2.3 | 900.0 |
| CTCL | HH | 40000 | 0.0 | 23.4 | 24000.0 |
| CTCL | HUT-78 | 900 | 13.5 | 0.8 | 1700.0 |
| PTCL | FEPD | 3 | 18.4 | 0.5 | 750.0 |
| Canine B cell lymphoma | CLBL1 | 225 | 0.0 | 0.5 | 175.0 |
| murine B cell lymphoma | A20 | 2450 | 9.5 | 1.0 | 850.0 |
| murine B cell lymphoma | BCL1clone5B1b | 500 | 0.0 | 0.5 | 435.0 |

**Figure S1. Camidanlumab tesirine *in vitro* activity in DLBCL cell lines is not affected by the presence of *BCL2* or *MYC* translocation or *TP53* inactivation.** Graphs show the distribution of camidanlumab tesirine (ADCT-301) IC50 values in DLBCL cell lines with or without *BCL2* translocation, *MYC* alterations and *TP53* inactivation.

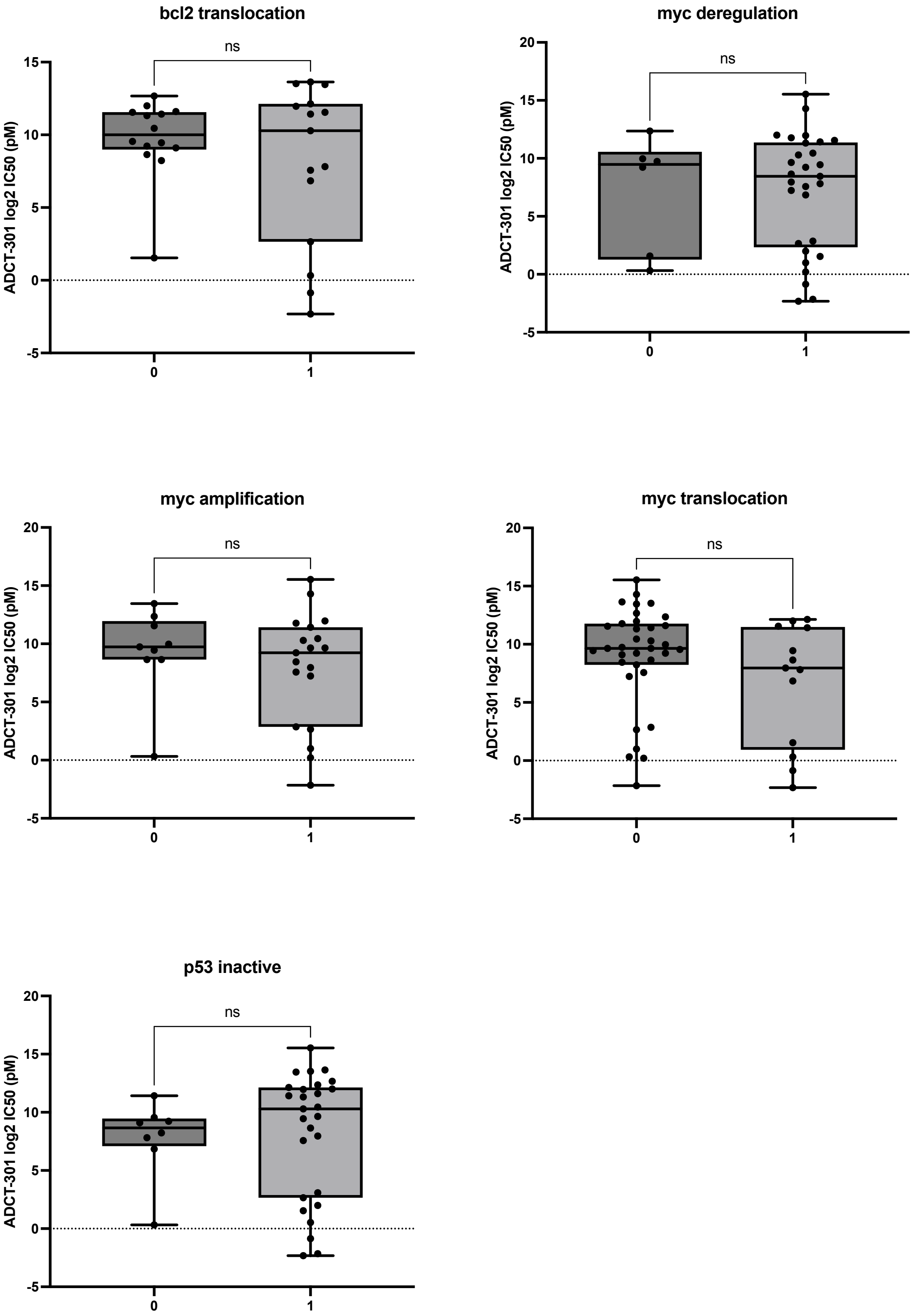
